# Learning a chemistry-aware latent space for molecular encoding and generation with a large-scale Transformer Variational Autoencoder

**DOI:** 10.64898/2025.12.19.695394

**Authors:** Hugo Talibart, Dimitri Gilis

## Abstract

Searching for molecules optimizing certain properties remains a key challenge due to the vastness of the chemical space, its discrete nature, and the limited availability of bioactivity data. One way to address these issues is to build a mapping from the chemical space to a continuous latent embedding space where efficient exploration and smooth interpolations become possible. Existing methods suffer from several limitations: failure to accurately reconstruct molecules, poorly structured latent space where whole regions cannot be decoded to valid molecular graphs or where distances fail to reflect chemical similarities, not to mention the unavailability of ready-to-use code or models trained on sufficiently large datasets, limiting their practical application. In this work, we provide a large-scale pre-trained Variational Autoencoder based on the Transformer architecture to convert small organic molecules to continuous fixed-size embeddings in a chemistry-aware structured latent space and back. With a **97%** reconstruction rate and a **100%** validity rate, the model provides a reliable mapping and is able to generate a high diversity of novel molecules. We introduce a training objective with a novel loss term explicitly enforcing embedding distances to reflect Tanimoto similarities between molecular fingerprints, leading to a smooth and well-structured latent space enabling seamless exploration and interpolation. We show that our model achieves solid performances on molecular optimization tasks compared to other generative models. We release an open-source, user-friendly implementation as a pip package and at github.com/3BioCompBio/chembed, along with pre-trained models, that can be readily used or fine-tuned for downstream drug design applications.

**Scientific contribution:** We introduce a large-scale SELFIES Transformer VAE with a novel loss aligning latent space distances with chemical similarity. This yields a highly structured, chemistry-aware latent space, alongside high reconstruction accuracy. Our open-source release with pre-trained weights makes the model directly usable and adaptable for molecular design applications.

## 1 Introduction

Drug design is a time-consuming and expensive process: it is estimated that developing a single drug may take 10 to 20 years and cost around 2 billion dollars [1, 2]. Candidate molecules must be identified within the vast chemical space, where the number of theoretically synthesizable drug-like compounds was estimated to be in the order of 10^60^ [3]. The virtually unlimited size and inherently discrete nature of this chemical space make it difficult to explore, to design molecules with desired features, targeting specific receptors or optimizing certain properties. Furthermore, in contrast with the vastness of the chemical space, the amount of available annotated data on molecular properties and bioactivity is astronomically small. Hence, while deep learning models have achieved spectacular breakthroughs in areas such as computer vision and natural language processing and their application to drug design related tasks is compelling, their effectiveness in real-life scenarios may be hindered by the scarcity and quality of available data for the task at hand. Training a model from scratch on each dataset risks overfitting, poor generalization, or low diversity among proposed molecules. In light of the scarcity of annotated data relative to the vastness of the search space, there is a need for unsupervised computational methods to make use of unlabelled molecular datasets to map the molecular space to a latent embedding space, to provide versatile representations for downstream tasks and enable smooth exploration. To these aims, the learned embeddings should capture essential and meaningful patterns in the original data, their structure should reflect their underlying relationships, and the mapping between original molecules and their embeddings should be reversible.

In the field of molecular design, different methods were proposed over the years to map the chemical space to a latent embedding space. Early work on this matter was mainly based on variational autoencoders [4] trained to reconstruct molecular graphs [5] or string representations such as SMILES [6]. In contrast with regular autoencoders, variational autoencoders learn a probabilistic distribution over the latent space, enabling sampling and interpolation by enforcing continuity and structure in the latent representations. The choice of which molecular representation to be reconstructed comes with trade-offs: graph-based representations may offer more faithful and unique representations of the molecular structures but are more computationally demanding, while string-based representations such as SMILES [7] are more efficient. As models may struggle to learn the specific syntax of SMILES and produce strings that cannot be translated to semantically valid molecules, potentially leading to large “dead areas” in latent spaces [6], SELFIES [8] emerged as a promising alternative, ensuring that all token sequences correspond to semantically valid molecular graphs, while retaining the efficiency of linear notation approaches. Representing molecules as token sequences made it possible to capitalize on deep learning advances in Natural Language Processing. In particular, following the success of Transformers [9], their architecture and training strategies [10] were soon adopted in the field of drug design for tasks such as property prediction [11–13] and molecular generation [14]. The attention mechanism in Transformers enables them to focus on relevant parts of the inputs and capture long-range dependencies between tokens, which is of particular relevance in the context of molecules, where tokens represent atoms, rings and branches that interact with each other. Despite the success of pretrained models and their generative capacities, their latent space lacks regularization and structure, making them less suitable for controlled molecular optimization and chemical space navigation, as they cannot produce fixed-size, self-contained embeddings that are directly decodable, nor do they support smooth interpolations and transitions in their latent space.

The question of latent space semantic consistency remains largely underexplored in the field of small molecules design. Most models rely at most on a Kullback-Leibler divergence loss term alone to regularize latent space structure. Although this strategy encourages some continuity between input representations, it does not enforce any semantic continuity between the initial objects that are represented. Indeed, two sequences of tokens may differ by only a few token edits but represent significantly different molecules, and conversely, two molecules that are closely related may be represented by significantly different linear notations, as SMILES or SELFIES canonicalization only addresses the question of unicity. A few strategies were proposed to actively steer models to build a semantically meaningful latent space. Some strategies common in Bayesian Optimization consist in jointly training models with predictive tasks and/or periodically retrain them with new samples evaluated by the black-box objective functions to be optimized during the process [15, 16]. However, on top of a risk of overfitting [17], models may end up embedding linearly the target properties into few dimensions of the latent vectors [18]. Alternatively, a handful of methods have begun incorporating ideas from metric learning, a concept introduced in [19] which aims at learning mappings where distances in latent space are consistent with input class labels, and which was shown to improve generalization in predictive tasks. Grosnit et al. [20] introduced a molecular VAE trained with a triplet loss to enforce a discriminative latent space where “positive” and “negative” samples according to a threshold in the objective function to be optimized are clustered separately. A similar idea was also proposed by Koge et al. [18], with a log-ratio loss. More recently, Sellner et al. [21] trained SMILES Transformers with an additional loss term *L*(*A, X*) defined, for one anchor molecule *A* and all other samples *X* in the batch, as:

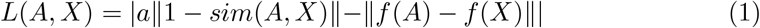

where *a* is a hyperparameter, *f* is the encoding function – here defined as a Transformer Encoder followed by a pooling operation – and *sim* can be any similarity measure between two molecules, which in practice they set to the Tanimoto similarity between molecular fingerprints [22]. Their method avoids the need to define positive and negative samples and showed substantial search space reduction in virtual screening tasks. However, embeddings obtained in this way cannot be decoded back to the original molecules, preventing their use in molecular optimization tasks.

In this paper, we introduce a SELFIES variational autoencoder with a Transformer-derived architecture trained on a large molecular dataset with a novel training objective focusing on reconstruction and latent space semantic consistency. This architecture leverages both the attention mechanism of Transformers and the capacity of the variational autoencoder framework to produce a smooth latent space. We address latent space structure with an auxiliary molecular prediction task and a novel loss term which explicitly enforces Euclidean distances between embeddings in latent space to be consistent with Tanimoto similarities between molecular fingerprints of encoded molecules. We show that this loss term induces a highly structured latent space that allows for smooth interpolations and exploration. The use of SELFIES representations enables computational efficiency and ensures that all vectors in latent space can be decoded to semantically valid molecular structures. Aware of the reconstruction issues reported in several widely cited models [23], we reduce the regularization as recommended in [24] to obtain high reconstruction rates. We show that, even with a naive search strategy, the model is able to propose relevant molecule candidates in molecular optimization tasks. Beyond the mere proof-of-concept, we prioritize usability by releasing a user-friendly implementation with complete code, environment and pre-trained models. We trained it on a large, stereochemistry-aware dataset derived from Pub-Chem [25] with a focus on small organic molecules within the typical bioactive range. The model provided is ready to use or fine-tune for tasks like molecular generation, optimization, and representation learning, sparing users the time and computational cost to train a large model from scratch.

## 2 Methods

This section outlines the key components of our approach. First, we describe how molecular structures are converted into standardized token sequences to provide as input to our model. Then, we detail the architecture of our model, which embeds the Transformer architecture into a variational framework to leverage attention mechanisms to map molecules to meaningful fixed-size latent vectors, enabling both reconstruction and generation of novel molecules. To ensure that this mapping is reliable and relevant for practical and efficient chemical space exploration, we designed a composite training objective which emphasizes reconstruction and latent space organization thanks to an auxiliary molecular prediction task and a novel loss term explicitly aligning latent embedding distances to Tanimoto similarities between molecular fingerprints. We specify the pre-processing steps used to construct the large dataset of small organic molecules on which our model was trained, and how token imbalance was handled. Then, we describe the simple algorithm for molecular optimization search used to validate our model. An overview of the model is depicted figure 1. Comprehensive implementation details including the full architecture, training setup and hyperparameters are provided in Additional file 1, section 1.

**Fig. 1.**
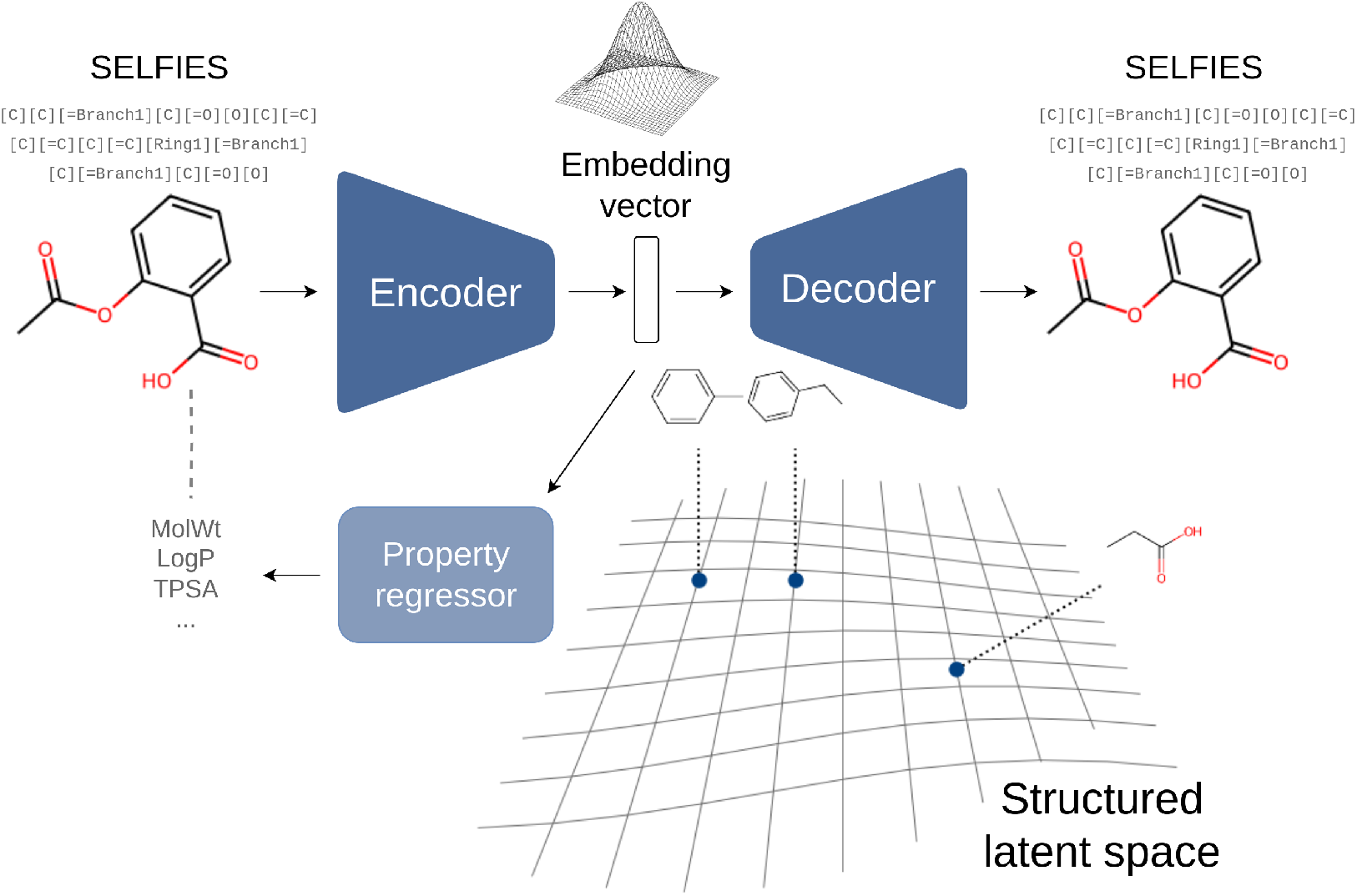
Overview of the model. Molecules are represented by SELFIES strings and fed to the encoder, which outputs an embedding vector following a Gaussian distribution. The decoder is able to decode any embedding vector into a SELFIES string, which in turn can be decoded to a valid molecular graph. A property regressor is jointly trained to predict chosen properties of the initial molecules from their latent embeddings alone, thereby encouraging more meaningful embeddings. By integrating an additional loss term explicitly aligning embedding distances to chemical similarities into the variational framework (see section 2.3), the latent space is structured so that neighboring vectors represent similar molecules.

### 2.1 Molecular representations

To avoid the computational cost associated with molecular graph matching which would make it prohibitively expensive to train it on large datasets, we adopted string-based representations. Since SMILES [7] representations often lead to large “dead areas” in latent space [6] as models struggle to learn the specific SMILES syntax, we used SELFIES representations instead [8], ensuring that all vectors in latent space can be decoded to semantically valid molecules. To ensure canonicity, molecules are standardized using the ChEMBL standardization procedure described in Bento et al. [26] and RDKit [27] before conversion to SELFIES. These SELFIES strings are tokenized using a vocabulary derived from the large dataset described in section 2.4. Though this vocabulary is large, occasionally some SELFIES strings may contain unseen tokens, in which case they may be replaced with the semantically closest tokens according to a specifically crafted distance metric. Details on how this distance metric is defined are provided in Additional file 1, section 2.

### 2.2 Autoencoder architecture

The autoencoder consists in two modules: an encoder that maps SELFIES tokens to latent vectors of dimension (1, *d*) with *d* = 256 and a decoder that maps latent vectors back to SELFIES tokens. Both modules are based on the Transformer architecture [9], allowing them to capture long-range interactions between tokens. Tokens are embedded via an embedding layer initialized using an MDS projection based on semantic distances (as described in section 2.5), combined with positional encoding, before being fed to a Transformer encoder stacking 6 layers with multi-head attention and feed-forward. To embed the Transformer into a variational autoencoder framework [4], a bottleneck is introduced by using only the first position of the Transformer encoder output to derive mean and standard deviation vectors parameterizing the Gaussian distribution from which the latent vector is then sampled (see figure 2). To decode a latent vector, it is provided as memory input to a 6-layer Transformer decoder. During training (figure 3), the decoder inputs the target tokens shifted by one position, while at inference time (figure 4), it autoregressively generates tokens by conditioning on previously decoded outputs using a greedy decoding strategy. Target tokens are embedded the same way as in the encoder module, and the output logits, which represent unnormalized likelihood scores for each token in the vocabulary, are produced by applying the transpose of the embedding layer.

**Fig. 2.**
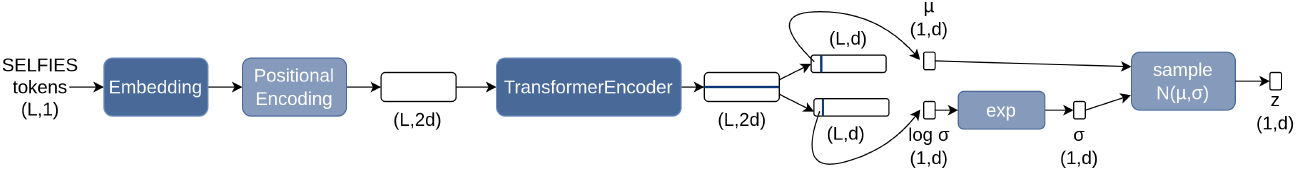
Encoder workflow at training time. The sequence of *L* SELFIES tokens is fed to an Embedding layer thoughtfully initialized and a Positional Encoding module to obtain a (*L*, 2*d*) tensor which is then fed to a Transformer Encoder module consisting of 6 stacked Transformer Encoder layers with multi-head attention and feedforward. The output of the Transformer Encoder is split horizontally and only the first position of each vector is retained to derive the mean *µ* and standard deviation *σ* defining the Gaussian distribution from which the latent embedding vector *z* is sampled. At inference time, the encoder is deterministic and only the mean is used.

**Fig. 3.**
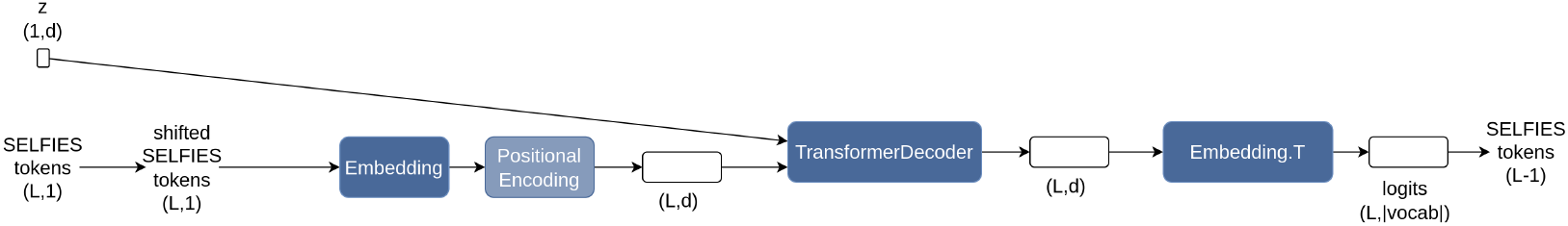
Decoder workflow at training time. Target tokens are shifted and fed to an Embedding layer and a Positional Encoding module in the same way as in the Encoder to obtain a (*L, d*) tensor which is fed to a Transformer Decoder module consisting of 6 stacked Transformer Decoder layers along with the embedding vector *z*. The output of the Transformer Decoder is converted to logits by applying the transpose of the input Embedding layer, defining a distribution over the SELFIES token vocabulary from which tokens can be sampled.

**Fig. 4.**
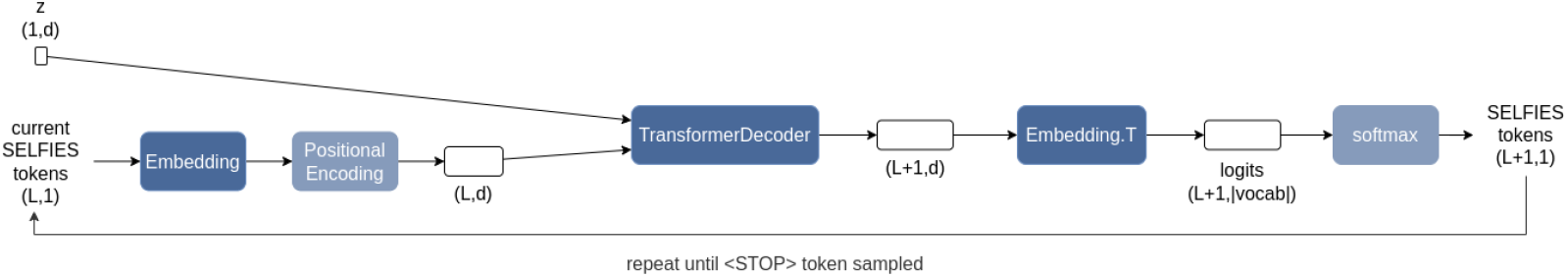
Decoder workflow at inference time. The decoder operates similarly to training, except that instead of receiving target tokens it generates them sequentially in a greedy fashion until a “STOP” token is sampled.

### 2.3 Training objective

The training objective seeks to balance accurate reconstruction of input molecules with the enforcement of a smooth and semantically meaningful latent space. To this end, we include a reconstruction loss, ℒ_*rec*_, penalizing discrepancies between the input and the output of the decoder, defined for a single sample of length *L* as the cross-entropy between one-hot encoded input tokens *y* and the predicted token distributions ŷ:

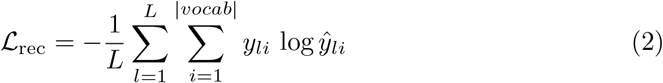

Acting as a regularizer, a Kullback-Leibler (KL) divergence loss term, ℒ_*KL*_, pushes the latent embeddings distribution towards a normal prior:

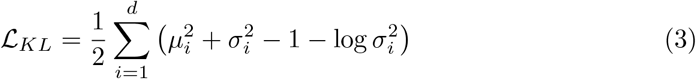

where *µ* and *σ* are respectively the mean and standard deviation of the distribution from which the latent vector is sampled.

While the KL divergence alone may be sufficient in certain datasets to provide an acceptable smoothness and structure of latent space, it should be noted that representing molecules with strings such as SELFIES implies an additional layer of abstraction between the target molecules and their representations. Indeed, two sequences of SELFIES tokens may differ by only a few token edits but represent significantly different molecules, particularly when substitutions affect topological tokens. Conversely, despite efforts towards canonicalization, very similar molecules may sometimes be represented by dissimilar SELFIES strings. To address this and improve latent space structure, we introduce a novel loss term, termed *Tanimoto loss*, to explicitly enforce Euclidean distances between vectors in latent space to be consistent with similarities between the molecules they represent, measured as the Tanimoto similarities [22] between Morgan fingerprints [28] computed by RDKit (with radius 2 and 1024 bits). Given a batch of *N* molecules, let *T* denote the *N* × *N* Tanimoto similarity matrix where *T*_*ij*_ = *Tanimoto*(*mol*_*i*_, *mol*_*j*_) and *D* denote the *N* × *N* Euclidean distance matrix where *D*_*ij*_ = ∥ *z*_*i*_ − *z*_*j*_ ∥ where *z*_*i*_ is the latent embedding of molecule *i*, then the Tanimoto loss ℒ_*Tanimoto*_ is defined as the correlation between the upper triangular values of the two matrices, shifted by 1 to be positive :

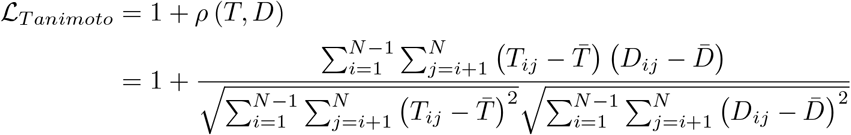

where 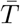 and 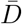 are the means over the upper-triangular entries.

Finally, to further improve the semantic meaningfulness of latent embeddings, we incorporate an auxiliary molecular prediction task alongside the autoencoding objective by jointly training a regressor – a simple multi-layer perceptron with two hidden layers with respectively 32 and 16 neurons and ReLU activations – to predict selected properties from latent vectors. By default, predicted properties include molecular weight, logP and polar surface area, as well as four graph topological descriptors: the Bertz CT index [29] which reflects molecular complexity, and the Kappa 1, 2, 3 indexes [30] which capture, respectively, the “cyclicity” of molecules, the “spatial density of atoms” and the “centrality of branching”, leading to a total of *N*_*prop*_ = 7 properties. The regressor is trained to minimize the average mean squared error (MSE), ℒ_*prop*_, between its output 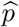 and (normalized) target properties *p*:

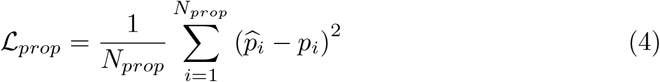

The complete loss function, ℒ, is defined as:

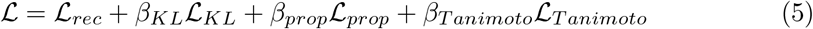

While the standard variational framework assigns the same weight to reconstruction and KL divergence loss terms, Yang et al. [24] showed that, in the context of molecular sequence autoencoders, the reconstruction loss is often underestimated. Following their recommendations, we set the KL divergence loss term coefficient *β*_*KL*_ to a lower value – in practice 0.2 – to improve reconstruction rates. We set *β*_*prop*_ = 1, *β*_*T animoto*_ = 0.5, as these values seemed to lead to better results in preliminary experiments.

### 2.4 Dataset and pre-processing

The model was pre-trained on a large dataset derived from PubChem [25]. Since our focus is on small organic molecules, all molecules with weight exceeding 600 Da, lacking carbon, or containing atoms not in the standard organic subset (B, Br, C, Cl, F, H, I, N, O, P, S, Si, Sn) were discarded, as well as molecules that could not be parsed by RDKit. The 600 Da cutoff was chosen based on Lipinski’s rule of five, which limits drug-like molecules to about 500 Da, extended with a margin to 600 to include heavier yet still bioactive molecules, consistent with the distribution of ligand weights in the PDB [31] and allowing for a very wide variety of compounds. This filtering led to a total of more than 95 million molecules. Salts and isotope values were removed and molecules were standardized as described in section 2.1. Note that stereochemistry information was kept, thereby increasing vocabulary size and model complexity but making it possible to handle stereochemistry-sensitive tasks. To train the property regressor, molecular properties were computed by RDKit (MolWt, logP, TPSA, BertzCT, Kappa1, Kappa2, Kappa3). The dataset was randomly split into train (80%) and test (20%) after removing any redundancy. Since Kappa2 and Kappa3 indexes may take extreme values, their values were clipped to their 0.001 and 0.999 quantiles before normalization on the train set. The train set was further split into an actual train set (80%) and a validation set (20%). Oversampling was applied to the train set as described in the next section.

### 2.5 Token imbalance handling strategies

In typical datasets of organic molecules, token occurrences are significantly imbalanced: in the dataset described in the previous section, `[C]` represents around 39% of all tokens whereas other tokens such as [/Si@@] appear only once. This poses significant challenges for the model, which may not be able to accurately reconstruct such rare tokens. To tackle these imbalances, we implemented four concomitant strategies: scaling gradients of embedding layers by inverse token frequencies, disabling weight decay in such layers, initializing them to reflect token semantic proximity, and upsampling SELFIES strings containing rare tokens. To initialize embedding layers, we defined an arbitrary distance metric between SELFIES tokens (formally defined in Additional file 1, section 2) and used scikit-learn’s implementation of the MDS algorithm to obtain token embeddings matching this distance matrix. As for the upsampling, each SELFIES string *S* = *S*_1_, · · ·, *S*_*L*_ in the dataset was assigned a *rarity score* defined as 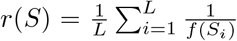 where *f*(*S*_*i*_) is the frequency of token *S*_*i*_ in the dataset, then each string was replicated *N* (*S*) = ⌈ *a × r*(*S*) ⌉ times, with *a* set to 10^−5^. This resulted in only a modest increase in the total number of training samples (76 886 404 vs 76 478 017), while ensuring that every token appeared at least 4 326 times.

### 2.6 Molecular optimization search algorithm

To evaluate the relevance of our autoencoder on molecular optimization tasks, we implemented a simple search strategy inspired by genetic algorithms. Given a fitness function *f* to be maximized and an initial set of *M* molecules, the algorithms proceeds as follows:

1. **Encoding**: The *M* molecules are encoded into seed latent vectors *z*_1_, ··,*z*_*M*_
2. **Crossover**: All pairwise interpolations between seed vectors are computed to form 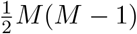 mixture vectors: 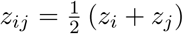
3. **Mutation**: Gaussian noise ~ 𝒩 (0, *σ*^2^) is added to all seed and mixture vectors to generate *N*_*pop*_ candidates (in practice we use *σ* = 0.5)
4. **Decoding**: All latent vectors are decoded into molecules
5. **Filtering**: Duplicates are removed, and steps 3–4 are repeated until a total population of *N*_*pop*_ unique molecules 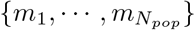 is sampled
6. **Evaluation**: The fitness function *f* to be maximized is evaluated on all molecules
7. **Selection**: Molecules *m*_*i*_ satisfying 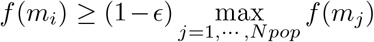 are retained as the next generation (in practice we use *ϵ* = 0.01)
8. Steps 1 – 7 are repeated *N*_*gen*_ times

## 3 Results and discussion

### 3.1 Reconstruction and generation

After training, the model was able to fully reconstruct 18 552 099 SELFIES strings out of the 19 119 505 SELFIES strings in the test set, i.e. a string reconstruction rate of 97%. The token reconstruction accuracy was 99.6%. This demonstrates strong fidelity of the autoencoder, with very few molecules failing to be reconstructed correctly. Out of the 567 406 output SELFIES that were not identical to the input SELFIES, 29 079 corresponded to identical molecules up to stereochemistry, and 37 378 corresponded to molecules with Tanimoto similarity of 1.0 with the initial molecule. This shows that the model captures both local syntax and global structure.

To evaluate the model’s generative capacities, 100 000 SELFIES strings were generated by feeding the decoder random samples from 𝒩 (0, 1). All generated samples were successfully decoded to valid molecular graphs by RDKit, yielding a validity rate of 100%. This means that our model does not suffer from dead regions in latent space. Almost all generated SELFIES (99.997%) were unique, with high internal diversity: *IntDiv*1 = 0.8848, *IntDiv*2 = 0.8813, where *IntDiv*1 and *IntDiv*2 are computed for a generated set *G* as in MOSES [32]:

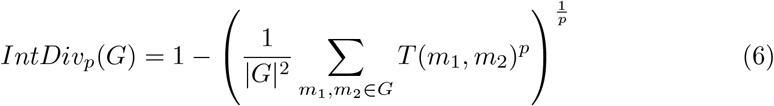

where *T* is the Tanimoto similarity. These high internal diversity values suggest that the model is able to explore broad regions of the chemical space. Only 29 of the 100 000 generated molecules were already in the train set, which corresponds to a novelty score of 99.97%. This score confirms that, while the model has excellent reconstruction performances, it did not simply memorize the training set and is able to extrapolate to novel molecules. Interestingly, 6 of the generated molecules were in the test set.

Since the goal of this large pre-trained model is to map a large portion of the chemical space rather than reproduce the properties of a small drug-like subset, we did not compute other MOSES metrics such as the fraction of molecules passing filters or similarity measures with a target set, as they are not informative in this context.

Together, these results indicate that the model is a reliable autoencoder, capable of faithfully reconstructing input molecules, and that it can also generate diverse molecules beyond the training set, supporting its potential for latent space exploration.

### 3.2 Latent space structure

#### Distance consistency analysis

To systematically assess whether the latent space effectively preserves chemical similarities, we sampled 1 000 000 random molecule pairs from the test set and computed the Tanimoto similarity between their Morgan fingerprints and the Euclidean distance between their embeddings. Results are plotted figure 5. The Pearson correlation is − 0.84, indicating a strong inverse relationship between distances in the latent space and similarities in the original chemical space. This confirms that the mapping successfully preserves global chemical similarity structure, making the latent space suitable for downstream local searches.

**Fig. 5.**
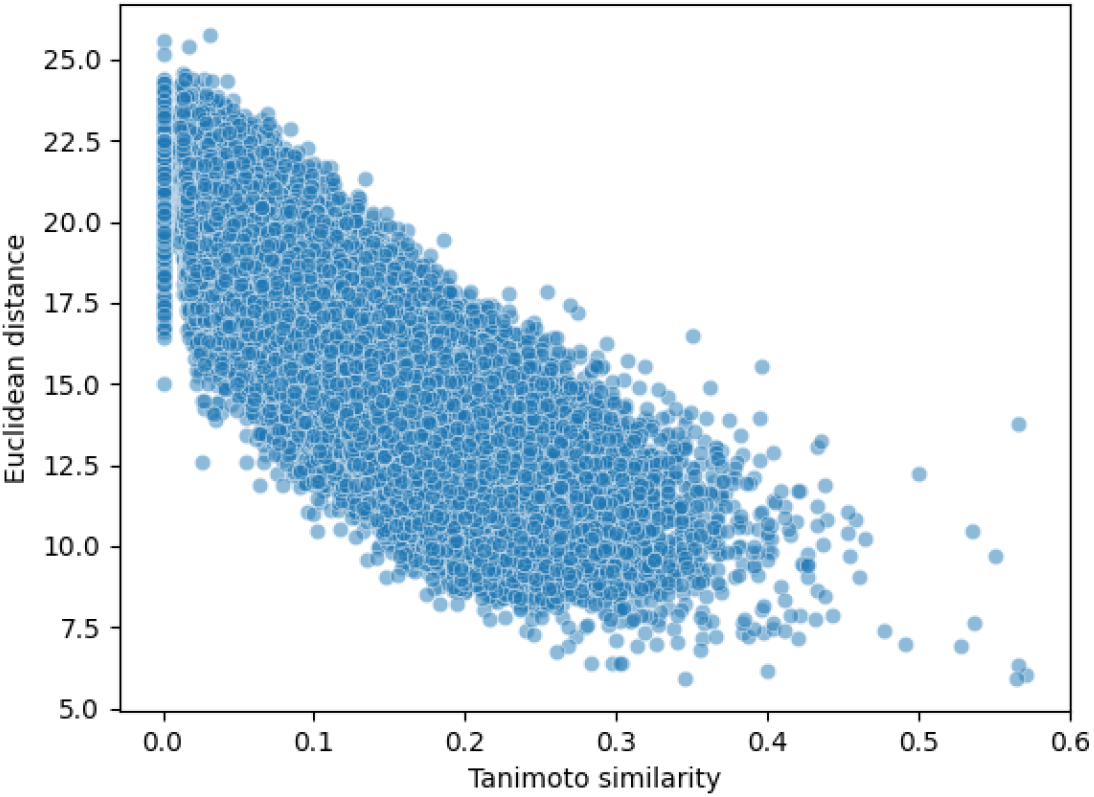
Chemical distance consistency visualization. Euclidean distance between embeddings versus Tanimoto similarity between Morgan fingerprints of initial molecules for 1 000 000 random molecule pairs from the test set.

#### Grid visualization

To qualitatively explore the latent space of our model, we examined the neighborhood of selected molecules with a random 2D grid visualization, following the approach of Kusner et al. [33]. A reference molecule was encoded into an embedding vector and two random orthonormal vectors were generated using the Gram-Schmidt process to define a 2D grid with the reference molecule at the center. Each point along the grid was decoded into a molecule, and its Tanimoto similarity with the reference molecule was computed in order to quantify how latent space shifts translate to structural changes. Figure 6 shows such a visualization with lidocaine at two different resolutions. In the local neighborhood (figures 6a and 6b), decoded molecules remain structurally similar to the reference, with Tanimoto similarities gradually decaying with distance, demonstrating local continuity. Expanding the grid to a wider neighborhood (figures 6c and 6d) reveals that, beyond the nearest neighborhood where molecules typically display high Tanimoto similarity with the reference (ranging from 0.6 to 1.0), the maximum attainable Tanimoto similarity drops sharply before leveling off, producing the wedge-shaped distribution in figure 6d. This pattern reflects the asymmetry that there are more ways for molecules to differ than to remain similar. The local geometry of the latent space is anisotropic: some directions may change the molecule only a little, while others induce drastic modifications in the scaffold. This is expected given the discrete nature of the chemical space and the many-to-one decoding. These local patterns are not visible in global pair analyses (figure 5), as they average over many references and directions, yielding a global linear trend. Overall, the grid visualizations show that the latent space is anisotropic and heteroscedastic, yet locally continuous. To the extent permitted by the discreteness of the chemical space, small shifts in latent space produce small structural changes.

**Fig. 6.**
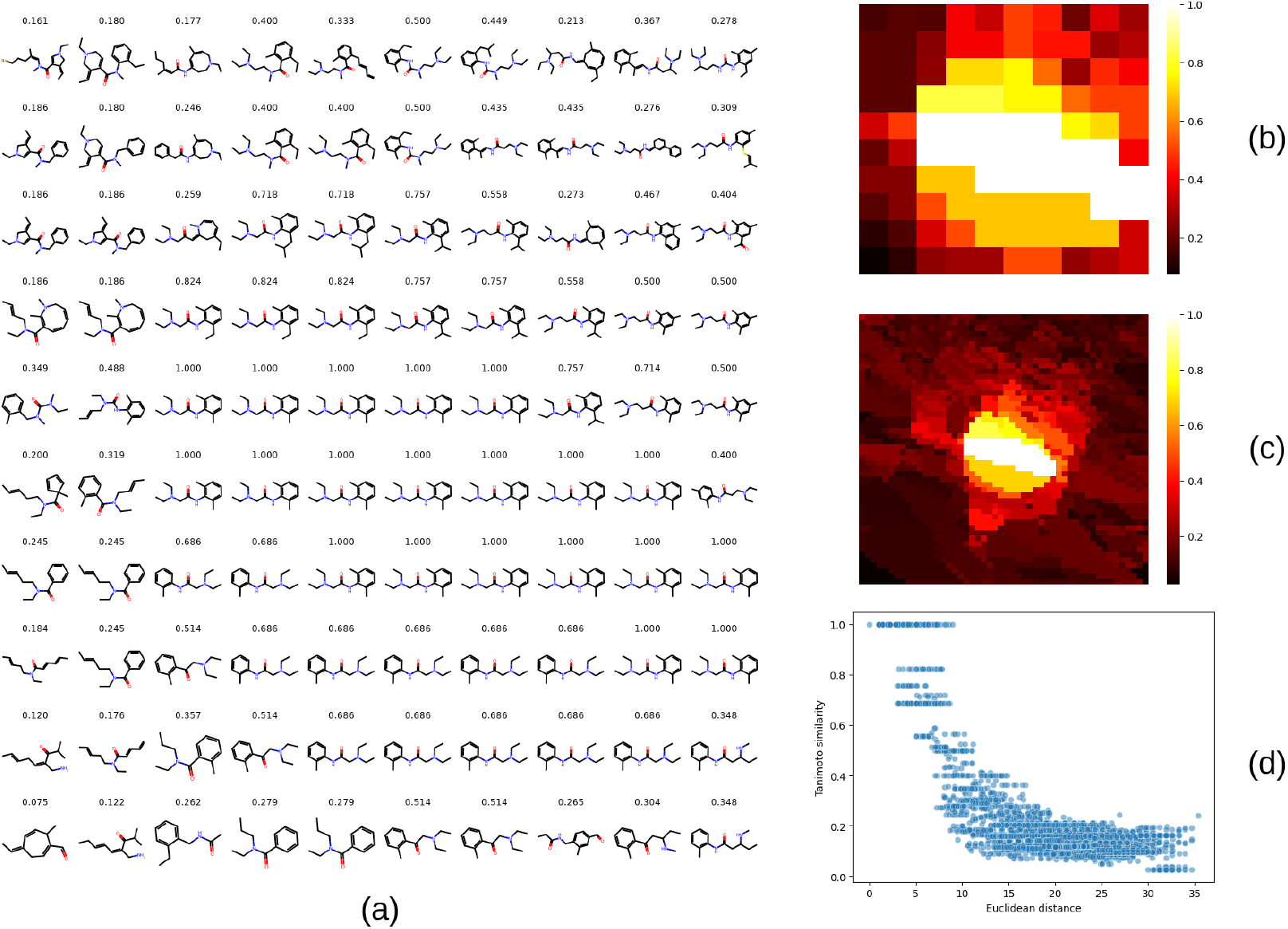
Latent space neighborhood of lidocaine. (a) Molecules decoded on a 10 *×* 10 grid using two orthonormal unit vectors reused across panels. Numbers above molecules indicate Tanimoto similarities with lidocaine. (b) Heatmap of panel (a)’s Tanimoto similarities. (c) Same heatmap on a larger 50*×* 50 grid with the same vectors. (d) For the 50 *×* 50 grid in (c), scatter plot of Tanimoto similarities versus Euclidean distances from the reference molecule’s embedding.

### 3.3 Tanimoto loss ablation study

To assess the contribution of the Tanimoto loss on latent space structure, we performed an ablation study. We sampled a representative subset of 1 000 000 molecules from the training set using stratified sampling, further split into train (80%) and validation (20%), and trained models for 5 epochs with the same architecture and the same hyperparameters as our main model, except without the auxiliary properties regression task. The Tanimoto loss coefficient was varied between 0 (no Tanimoto loss) and 1. After training, we measured the token reconstruction accuracy on the validation set, the Pearson correlation between Euclidean distances in latent space and Tanimoto similarities between initial molecules, and the trustworthiness of the latent space [34] – a score between 0 and 1 quantifying how well local neighborhoods are preserved. Each experiment was ran with 3 different seeds for robustness. As shown in figure 7, token reconstruction accuracy remains stable around 0.74, indicating that the Tanimoto loss does not harm reconstruction. On the other hand, as the Tanimoto loss coefficient increases, the Pearson correlation between Euclidean distances and Tanimoto similarities strengthens, from ≃ − 0.37 to ≃ − 0.79, and the trustworthiness increases from ≃ 0.77 to ≃ 0.93. These results indicate consistent improvements in both global and local latent space structures. Details on the ablation study methodology and additional results can be found in Additional file 1, section 3.

**Fig. 7.**
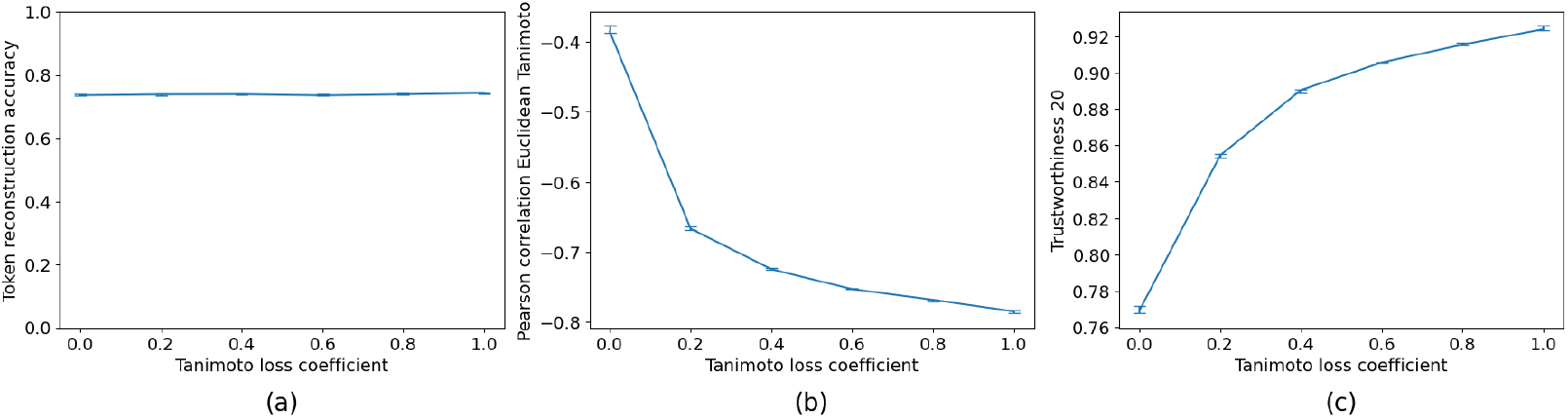
Impact of Tanimoto loss coefficient. (a) Token reconstruction accuracy, (b) Pearson correlation between Euclidean distances and Tanimoto similarities, and (c) trustworthiness (*k* = 20). Each curve shows the median across 3 seeds, with error bars indicating the range.

### 3.4 Molecular optimization

#### Benchmark overview

To demonstrate the relevance of a structured latent space in molecular optimization tasks, we evaluated the performances of our model combined with the simple search algorithm proposed in section 2.6 on the Tartarus benchmark [35]. This benchmark includes diverse tasks simulating real-life molecular design problems. For each task, fitness evaluation functions are provided as black-box workflows inputting SMILES strings and outputting fitness scores, utilizing semi-empirical programs based on well-established computational chemistry simulation methods. The goal for each task is to propose molecules optimizing the associated fitness function, with a maximum of 5 000 function evaluations allowed. Datasets with SMILES strings and associated precomputed fitness scores are also provided, enabling model fine-tuning and initialization of the optimization algorithm from the highest-scoring molecules. We discarded the chemical reaction substrates design task due to inconsistencies in its fitness evaluation function (outputs varied significantly between repeated evaluations of the same molecule), retaining three main tasks: design of organic photovoltaics, design of organic emitters, and design of protein ligands. Additional figures and detailed results including the molecules generated by our model in each task along with their fitness values, synthetic accessibility scores, and Tanimoto similarities to the initial molecules, as well as the best performing molecules from the initial datasets, are provided in Additional file 1, section 4.

#### Experimental setup

For each task, we used the provided dataset to fine-tune our pre-trained model by processing it as in section 2.4, randomly splitting it into a train set (80%) and a validation set (20%), making sure that all SELFIES tokens of the dataset appeared in the train set. All models were fine-tuned with a learning rate of 10^−5^ for 200 epochs, embeddings of tokens that were not in the pre-training dataset were initialized as described in section 2.5. For each subtask, candidate molecules were then obtained by applying the algorithm described in section 2.6 using the fine-tuned model, starting with the set of all molecules sharing the highest fitness score in the initial dataset (with a maximum of *M* = 20 initial molecules), setting the generation standard deviation parameter *σ* to 0.5 and the relative selection tolerance *ϵ* to 0.01. To respect the benchmark’s constraints, a maximum of *N*_*gen*_ = 10 iterations were performed with a population of *N*_*pop*_ = 500. Results are reported as means and standard deviations over 5 runs and are compared with the eight baseline methods from the original publication (SMILES-VAE [6], and its SELFIES reimplementation termed SELFIES-VAE, MoFlow [36], SMILES-LSTM-HC [37, 38] and its SELFIES reimplementation termed SELFIES-LSTM-HC, REINVENT [39] GB-GA [40] and JANUS [41]).

#### Design of Organic Photovoltaics

The organic photovoltaics design task consists in two subtasks which both aim at finding a molecule to be used in bulk heterojunction devices, with a synthethic accessibility constraint. In the first subtask, the molecule is to be used with 6,6-phenyl-C61-butyric acid methyl ester (“PCBM”) as acceptor, and in the second subtask is to be used with a poly[N90-heptadecanyl-2,7-carbazole-alt-5,5-(40,70-di-2-thienyl-20,10,30-benzothiadiazole)] (“PCDTBT”) as donor. The two fitness functions are multi-objective, combining power conversion efficiency (*PCE*) predicted using XTB [42] and CREST [43] with a synthetic accessibility score (*SA*_*score*_) predicted by RDKit [27]. Results are reported in table 1.

**Table 1.** Results for the “organic photovoltaic” optimization tasks. Comparaison of the results obtained with chembed, our model, with results of baseline models on two OPV tasks: *PCE*_*P CBM*_ − *SA*_*score*_ and *PCE*_*P CDT BT*_ − *SA*_*score*_. ↑ indicates higher is better, indicates lower is better. ↓ Values are mean *±* standard deviations over 5 runs. The best score in each columnn is shown in bold. The first row gives the score of the best molecule in the initial dataset.

| | $PCE_{PCBM} - SA_{score} \uparrow$ | $PCE_{PCDTBT} - SA_{score} \uparrow$ |
| --- | --- | --- |
| DATASET | 7.57 | 31.71 |
| **CHEMBED** | **$8.46 \pm 0.54$** | $28.04 \pm 0.99$ |
| SMILES-VAE | $7.44 \pm 0.28$ | $10.23 \pm 11.14$ |
| SELFIES-VAE | $7.05 \pm 0.66$ | $29.24 \pm 0.65$ |
| MoFlow | $7.08 \pm 0.31$ | $29.81 \pm 0.37$ |
| SMILES-LSTM-HC | $6.69 \pm 0.40$ | **$31.79 \pm 0.15$** |
| SELFIES-LSTM-HC | $7.40 \pm 0.41$ | $30.71 \pm 1.20$ |
| REINVENT | $7.48 \pm 0.11$ | $30.47 \pm 0.44$ |
| GB-GA | $7.78 \pm 0.02$ | $30.24 \pm 0.80$ |
| JANUS | $7.59 \pm 0.14$ | $31.34 \pm 0.74$ |

Our model achieves the best mean fitness score in the *PCE*_*P CBM*_ − *SA*_*score*_ maximization task, significantly outperforming other baseline methods. Generated molecules have moderate to high synthetic accessibility scores (between 4.59 and 5.09), akin to the initial molecule (4.59), and relatively low Tanimoto similarities with the latter (between 0.10 and 0.19). In the *PCE*_*P CDT BT*_ − *SA*_*score*_ maximization task, like other baseline methods except for SMILES-LSTM-HC, our model fails to propose molecules with fitness scores higher than the best molecule in the initial dataset. Generated molecules vary greatly in their Tanimoto similarity to the initial molecule (between 0.18 and 0.55) and their synthetic accessibility scores (between 2.90 and 4.35).

#### Design of Organic Emitters

In the organic emitters task, the goal is to design emissive materials for organic light-emitting diodes (OLEDs) using thermally activated delayed fluorescence (TADF). The first subtask (“singlet-triplet”) consists in finding molecules with a minimal energy gap between their singlet and triplet states, corresponding to improved emission efficency, the second subtask (“oscillator strength”) consists in finding molecules with maximal oscillator strength between the first excited singlet and the ground state, corresponding to improved fluorescence rates, and the third subtask (“multi-objective”) is a multiobjective optimization combining the first two goals while also keeping the excitation energies within the range of blue light. The fitness evaluation workflow is based on pyscf [44], CREST [43] and XTB [42]. Results are reported in table 2.

**Table 2.** Results for the “organic emitters” optimization tasks. Comparaison of the results obtained with chembed, our model, with results of baseline models on three tasks: singlet-triplet minimization, oscillator strength maximization, multi-objective. Values are mean *±* standard deviations over 5 runs. ↑ indicates higher is better, ↓indicates lower is better. The best score in each columnn is shown in bold. The first row gives the score of the best molecule in the initial dataset

|  | Singlet-triplet ↓ | Oscillator strength ↑ | Multi-objective ↑ |
| --- | --- | --- | --- |
| Dataset | 0.020 | 2.97 | −0.04 |
| **CHEMBED** | **0.003 ± 0.002** | **3.40 ± 0.34** | −0.04 ± 0.0 |
| SMILES-VAE | 0.071 ± 0.003 | 0.50 ± 0.27 | −0.57 ± 0.33 |
| SELFIES-VAE | 0.016 ± 0.001 | 0.36 ± 0.31 | **0.17 ± 0.10** |
| MoFlow | 0.013 ± 0.001 | 0.81 ± 0.11 | −0.04 ± 0.06 |
| SMILES-LSTM-HC | 0.015 ± 0.002 | 1.00 ± 0.01 | −0.24 ± 0.01 |
| SELFIES-LSTM-HC | 0.013 ± 0.003 | 1.00 ± 0.01 | −0.24 ± 0.01 |
| REINVENT | 0.014 ± 0.003 | 1.16 ± 0.18 | −0.15 ± 0.05 |
| GB-GA | 0.012 ± 0.002 | 2.14 ± 0.45 | 0.07 ± 0.03 |
| JANUS | 0.008 ± 0.001 | 2.07 ± 0.16 | 0.02 ± 0.05 |

Our model significantly outperforms all other methods in the singlet-triplet minimization task. Generated molecules (figure S7 of Additional file 1) have moderate to low synthetic accessibilities (between 4.18 and 5.62) and their Tanimoto similarities with the initial molecules range from 0.09 to 0.27. One generated molecule in particular has a strikingly low singlet-triplet value (7 × 10^−15^). The model also outperforms other baseline methods in the oscillator strength maximization task, however only one novel unique molecule was proposed, in 2 out of the 5 runs, while in the other 3 runs it did not propose a better molecule than the initial one. In the multi-objective task, however, the model completely failed to propose better novel molecules in all 5 runs, as most baseline methods.

#### Design of Protein Ligands

In the protein ligands tasks, the goal is to identify molecules with minimal docking scores – indicating maximal predicted affinity – with a target of interest, as computed by the docking program QuickVina2 [45], with drug-likeliness and synthethic accessibility filters: molecules with Synthetic Accessibility score above 4.5 or QED scores below 0.3 (as computed by RDKit) are assigned a score of 10^4^. Three targets are considered, with PDB IDs respectively 1SYH, 4LDE and 6Y2F. Results are reported in table 3.

**Table 3.** Results for the protein ligands design task. Comparaison of the results obtained with chembed, our model, with results of baseline models on binding affinity maximization for three targets: 1SYH, 6Y2F and 4LDE. Values are mean *±* standard deviations over 5 runs. ↑indicates higher is better, ↓ indicates lower is better. The best score in each columnn is shown in bold. The first row gives the score of the best molecule in the initial dataset.

|  | 1SYH ↓ | 6Y2F ↓ | 4LDE ↓ |
| --- | --- | --- | --- |
| DATASET | −9.9 | −8.2 | −12.2 |
| **CHEMBED** | −10.9 ± 0.2 | −8.0 ± 0.4 | **−13.1 ± 1.0** |
| SMILES-VAE | −10.0 ± 0.7 | −8.3 ± 0.6 | −10.7 ± 0.2 |
| SELFIES-VAE | −10.4 ± 0.3 | −9.6 ± 0.5 | −11.1 ± 0.4 |
| MoFlow | −10.6 ± 0.4 | −9.8 ± 0.4 | −12.1 ± 0.4 |
| SMILES-LSTM-HC | −10.6 ± 0.6 | −10.0 ± 0.6 | −12.4 ± 0.3 |
| SELFIES-LSTM-HC | −10.8 ± 0.5 | −10.4 ± 0.4 | −12.7 ± 0.3 |
| REINVENT | **−11.8 ± 0.4** | −11.1 ± 0.3 | −12.8 ± 0.2 |
| GB-GA | −11.6 ± 0.5 | −10.9 ± 0.2 | −12.9 ± 0.1 |
| JANUS | −11.7 ± 0.4 | **−11.3 ± 0.3** | −12.8 ± 0.2 |

Our method’s performance on these docking optimization tasks is uneven. For target 4LDE, our method outperforms all other baseline methods, with an average docking score of output molecules of −13.11 kcal/mol. For target 1SYH, our performance is mid-range compared to other methods, with an average docking score of − 10.9 kcal/mol, which is better than the best molecule in the dataset (− 9.9 kcal/mol) but REINVENT, GB-GA and JANUS proposed molecules with significantly better docking scores (resp. −11.8, −11.6 and −11.7 kcal/mol). Tanimoto similarities of proposed molecules with the best initial molecules range from 0.18 to 0.25. However, for target 6Y2F, our method performs worst against compared methods, failing to propose a better molecule than the initial one. As shown in figures S11, S13 and S15 in section 4 of Additional file 1, molecules proposed by our search algorithm are (by design) close to initial molecules in the dataset. In particular, all molecules proposed for target 6Y2F contain a sulfonyl group, often bounded to a nitrogen, like the initial molecule. This occurs because this initial molecule is surrounded by a large region populated by molecules containing this group. Consequently, the algorithm is constrained to explore only molecules with this group. In contrast, best molecules generated by other baseline methods for this target do not contain this group, as shown in the original paper’s supplementary data. This example illustrates the main limitation of the search algorithm used here: its strong dependence on the initial molecule and its underlying assumption that molecular similarity correlates with fitness similarity.

#### Overall performances and discussion

Our model, when paired with a simple search strategy, demonstrates competitive performance across diverse molecular design tasks, achieving top performance in four out of the eight subtasks of the benchmark. This indicates that its latent space is sufficiently well structured to support efficient black-box function optimization, consistent with the continuity observed, where small shifts produce gradual property changes. However, in three subtasks, the search algorithm failed to identify molecules with improved fitness scores over the initial dataset. In two of these cases, the difficulty is shared by other baseline methods, reflecting the inherent challenge, with multiobjective optimization being particularly complex. Failures may reflect the anisotropy and heteroscedasticity previously observed, where some directions result in drastic scaffold changes. The third case, the 6Y2F docking task, highlights one core limitation of the search algorithm itself: it depends strongly on the initial molecule. One way to address this issue would be to, instead of using fixed generation standard deviation *σ* and relative selection tolerance *ϵ*, dynamically adapt these exploration parameters based on the local neighborhood structure to allow a broader exploration of the latent space when fitness stalls. The other core limitation is that it assumes that molecular similarity correlates with fitness, a premise frequently invalidated by activity cliffs [46]. It should be noted that no method consistently outperforms others across all tasks: while approaches such as REINVENT or JANUS perform better in some tasks, our model achieves the best results in others, showing its overall competitiveness despite the simplicity of its search strategy. That being said, it should be recognized that while the generated molecules can reach high fitness scores, many may be challenging to synthesize. This limitation directly arises from the intended purpose of our model, which is to map the entire chemical space rather than just a synthesizable or drug-like subset. As a result, many regions in the latent space decode to molecules that are chemically valid but not synthetically accessible or suitable drug candidates, reflecting the diversity and representativeness of the broader chemical space. Experiments reported in this section aimed primarily at demonstrating the utility of our model’s structured latent space for molecular optimization when combined with a simple search algorithm. However, stronger performances could likely be achieved with more advanced optimization strategies, such as reinforcement learning, the incorporation of explicit synthetic accessibility constraints, or adapting the search procedure to each specific task to mitigate current limitations. Taken together, these results indicate that, while the choice of search strategy is critical, the latent space is sufficiently structured to support competitive performance across diverse molecular optimization tasks.

## 4 Conclusions

In this paper, we introduced a large-scale Transformer VAE based on SELFIES to map small organic molecules to continuous fixed-size embeddings and back. The model was pretrained on a large dataset, providing broad coverage of the chemical space. It achieves high fidelity reconstruction (97%) and perfect validity (100%) in generation, providing a reliable mapping. High diversity and novelty of generated molecules indicate that it did not simply memorize the training set and is able to extrapolate to novel molecules. We introduced a training objective with a novel loss term, the Tanimoto loss, which explicitly aligns Euclidean distances between embeddings in the latent space to similarities in the chemical space. Analysis of the latent space revealed a strong inverse relationship and local continuity, indicating that it captures meaningful chemical structure, thereby providing a practical basis for molecular exploration and optimization. Experiments on the Tartarus molecular design benchmark showed competitive performance in four out of eight design tasks even with a simple search algorithm, while underperformance in others highlighted limitations more rooted in the search strategy than the model itself. Further improvements in the search algorithm that incorporate explicit constraints in synthetic accessibility, task-specific search and generally more advanced optimization methods, could lead to improvements in molecular design tasks. By releasing a ready-to-use large pre-trained model with an open-source user-friendly implementation, we aim to provide a foundation for the molecular machine learning community, to be built upon and leveraged for advances in downstream tasks such as molecular optimization, drug discovery and materials design.

## Abbreviations

KL: Kullback-Leibler
MDS: Multidimensional scaling
MSE: Mean Squared Error
ReLU: Rectified Linear Unit
VAE: Variational AutoEncoder

## Declarations

### Availability of data and materials

The software is available as a pip package (`pip install chembed`). All code, experiment scripts, example notebooks are available at https://github.com/3BioCompBio/chembed. The training and test datasets, derived from PubChem, are available on Zenodo at https://doi.org/10.5281/zenodo.17277040. All resources are released under the MIT License.

### Supplementary information

Additional file 1 contains the following supplementary materials:

- Detailed model architecture and implementation
- Definition of SELFIES token semantic distance
- Methodology and additional results of the ablation study
- For each task in the molecular optimization benchmark: best performing molecules from the initial dataset and molecules generated by chembed, with associated synthetic accessibility scores and Tanimoto similarities

### Authors’ contributions

HT and DG conceptualized the work. HT developed the methodology, implemented the software, carried out experimentations and drafted the initial manuscript. DG contributed to the manuscript. All authors read and approved the final manuscript.

## Acknowledgements

The present research benefited from computational resources made available on Lucia, the Tier-1 supercomputer of the Walloon Region, infrastructure funded by the Walloon Region under the grant agreement n°1910247.

