## Supplementary Materials for "Learning a chemistry-aware latent space for molecular encoding and generation with a large-scale Transformer Variational Autoencoder"

#### 1 Implementation details

Our implementation is based on PyTorch and PyTorch Lightning using Python 3.12. The maximum SELFIES length was set to 256. The encoder includes 6 Transformer encoder layers with multi-head self-attention (8 heads- and feed-forward networks (hidden dimension 1024) with ReLU activation, layer normalization and dropout (0.1). The latent embedding has dimensionality 256. The decoder includes 6 Transformer decoder layers with masked multi-head self-attention (8 heads), cross-attention and feed-forward networks (hidden dimension 512) with ReLU activation, layer normalization and dropout. In total, the autoencoder has 17.6 M Parameters. The regressor is a multilayer perceptron with two hidden layers (dimensions resp. 32 and 16) with ReLU activations, for a total of 8.9 K parameters.

The base model was trained with the Adam optimizer using a batch size of 256, a learning rate of $10^{-4}$ and a warm-up schedule of 100 steps. To improve training stability, gradient values were clipped to 1.0 to avoid exploding gradients, and the decoder and the regressor were trained only every 5 steps. Weight decay was applied to all layers except for the embedding layers with a rate of $10^{-5}$. Pre-training did not use a fixed number of epochs or early stopping: we monitored the validation loss and stopped when it reached a plateau, which occurred after approximately 3.41 M steps (22 epochs). Training was performed on a single NVIDIA RTX 5000 Ada GPU, using medium float32 precision to improve efficiency. For fine-tuning, models were trained with a learning rate of $10^{-5}$ for a fixed 200 epochs for simplicity.

#### 2 SELFIES tokens semantic distance

To initialize embedding layers and handle the encoding of tokens that are not in the initial vocabulary, we define a custom distance metric between SELFIES tokens that reflect their semantic meanings. Notice that each SELFIES tokens are follows a consistent format:

1. Opening bracket `[`

2. Bond indicator: one of `#` (triple), `=` (double), `-/`, `-\`, `/`, `\` or none (single)
3. Main symbol: an atomic element (e.g. `C`, `Br`) or a topology marker (e.g. `Branch1`, `Ring2`)
4. Chirality indicator: `@` or `@@` or none
5. Explicit hydrogen count: (`H`, `H0`, `H1`, etc. or none)
6. Charge indicator: `+1`, `-1`, `+2`, etc. or none
7. Closing bracket `]`

We define the distance between two tokens as a weighted sum of component-wise distances, each rescaled to be between 0 and 1:

- **Bond distance**: 0 if bond indicators match exactly, 0.1 if they differ only in stereochemistry but represent the same bond order, otherwise the average bond order. Weighted by 2.
- **Main symbol distance**: 0 if identical, 0.9 if both atoms or topological tokens, 1 otherwise. Weighted by 5.
- **Chirality distance**: 0 if chirality indicators match, 1 otherwise. Weighted by 1.
- **Hydrogen distance**: 1 if one token has no explicit hydrogen and the other one does, otherwise, the absolute difference in hydrogen count divided by 4. Weigthed by 1.
- **Charge distance**: Absolute charge difference divided by 5. Weighted by 1.

### 3 Ablation study

#### 3.1 Dataset construction

Because the original training set was too large for practical ablation studies, we derived a representative subset using stratified sampling. Molecules were stratified by SELFIES length ("short": $< \frac{1}{3}$ quantile $= 29$, "long": $> \frac{2}{3}$ quantile $= 45$, "medium": in between) and by presence/absence of rare tokens (defined as occurring $< 100\,000$ times in the training set), yielding 6 strata. A balanced sample of 1 000 000 molecules was drawn and split into training (80%) and validation (20%) sets. Oversampling was performed following the strategy described in section 2.5 of the main text.

#### 3.2 Experimental setup

We used the same architecture and hyperparameters as the main model, differing only in the training objective. Starting from a baseline with reconstruction loss and KL coefficient of 0.2, we varied the Tanimoto loss coefficient (0, 0.2, 0.4, 0.6, 0.8, 1.0), the KL coefficient (0, 0.2, 0.4, 0.6, 0.8, 1.0, 2.0), and the inclusion of an auxiliary property regression task. All experiments were ran for 5 epochs with 3 different random seeds.

##### 3.3 Evaluation metrics

After training, each model was evaluated on reconstruction, generation, and latent space structure: reconstruction on the full validation set, generation on 10 000 sampled molecules, and latent space structure on 2 000 validation molecules randomly sampled. Specifically, we measured:

- Token reconstruction accuracy
- KL divergence loss
- Uniqueness
- Internal diversity (IntDiv1, IntDiv2)
- SELFIES, molecule and fingerprint novelty scores
- Pearson correlation between Euclidean distances in latent space and Tanimoto similarities
- Pearson correlation between Euclidean distances in latent space and absolute difference of molecular properties
- Trustworthiness, defined in [1] as

$$T(k) = 1 - \frac{2}{nk\,(2n - 3k - 1)} \sum_{i=1}^{n} \sum_{j \in \mathcal{N}_i^k} \max\left(0, \left(r_i\left(j\right) - k\right)\right) \tag{1}$$

  We used $k = 5$, $k = 10$, $k = 20$ and $k = 30$.

##### 3.4 Additional results

As shown in figure S1, the KL divergence coefficient critically determines reconstruction quality. As it increases, token reconstruction accuracy decreases. For large coefficients, a posterior collapse occurs when the objective function contains only reconstruction and KL losses: the decoder outputs overly generic outputs that match tokens well on average but miss the finer structural distinctions between molecules, leading to a higher token reconstruction accuracy but a significant drop in the uniqueness of generated molecules. Including the property regression loss prevents this collapse, showing its role as regularizer. Furthermore, although the Tanimoto loss is the main driver of latent space organization, the property loss complements it by drawing molecules with similar properties closer together in latent space, as illustrated in figure S2.

Aside from the case of posterior collapse at high KL coefficients, generation diversity (uniqueness, IntDiv1, IntDiv2) and novelty (SELFIES, molecule, fingerprint) scores remained stable across experiments, showing no clear trend.

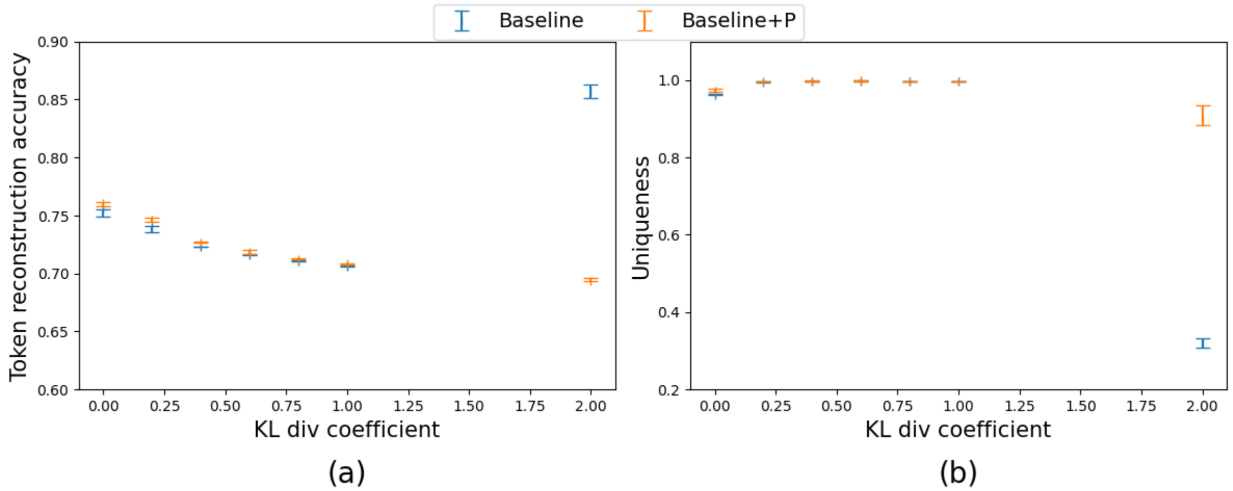


Figure S1: **Impact of the KL divergence loss coefficient.** (a) Token reconstruction accuracy versus KL divergence coefficient (b) Uniqueness of generated molecules versus KL divergence coefficient. Results are shown for the baseline (only reconstruction loss and KL divergence loss) in blue and for the baseline with the properties loss in orange. Each point shows the median across 3 seeds, with error bars indicating the range.

### 4 Molecular optimization additional results

#### 4.1 Design of Organic Photovoltaics

The best molecules proposed in each run for the organic photovoltaics design task are reported figure S3 for $PCE_{PCBM} - SA_{score}$ maximization and figure S5 for $PCE_{PCDTBT} - SA_{score}$ maximization, along with their fitness and synthetic accessibility scores and the Tanimoto similarity with the initial molecules (S4 and S6).

#### 4.2 Design of Organic Emitters

For the oscillator strength maximization task, the model outputted the starting molecule (figure S9) in 2 out of the 5 runs, and the molecule in figure S10 in the 3 other runs.

#### 4.3 Design of Protein Ligands

The initial dataset contains 1056 molecules with the best affinity with 4LDE ($-12.2$).

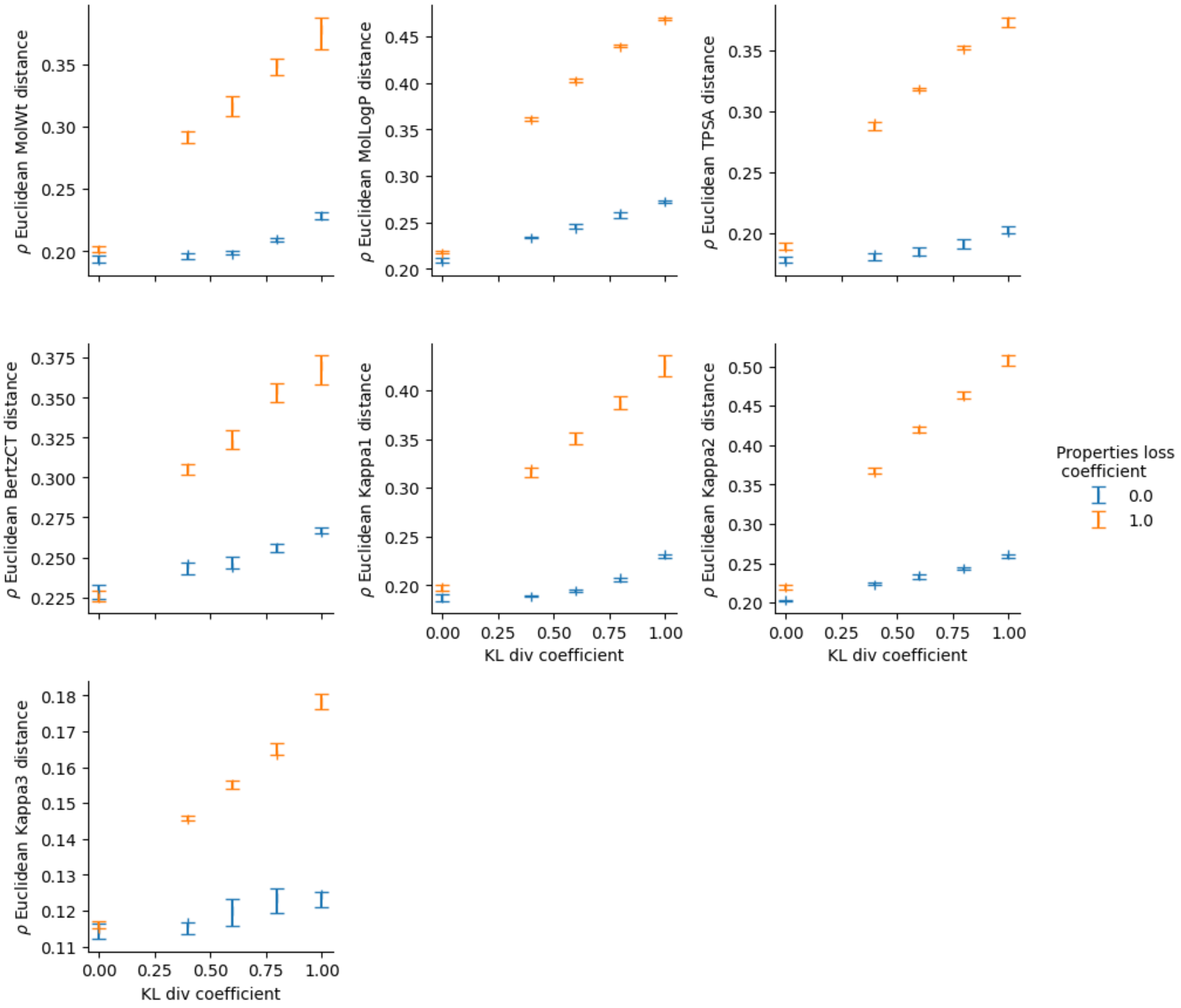


Figure S2: **Impact of the properties loss on latent space structure.** Pearson correlation between Euclidean distances in latent space and absolute value of differences in molecular property values in chemical space for all regressed properties (molecular weight, logP coefficient, polar surface area, Bertz CT and Kappa 1, Kappa 2, Kappa 3 indexes). Results are shown for the baseline the addition of the Tanimoto loss and the properties loss (in orange) and for the baseline with the Tanimoto loss only (in blue).

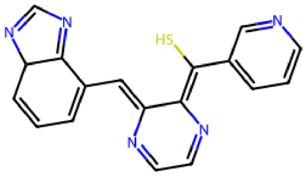

(a) Fitness: 8.40
$SA_{score}$: 4.59
Tanimoto: 0.13

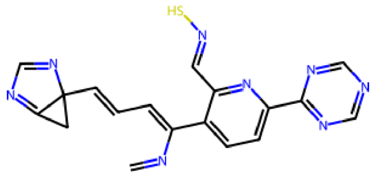

(b) Fitness: 7.58
$SA_{score}$: 5.14
Tanimoto: 0.19

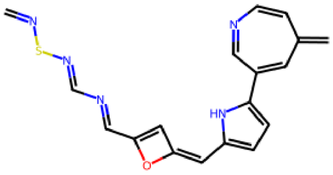

(c) Fitness: 8.27
$SA_{score}$: 5.08
Tanimoto: 0.10

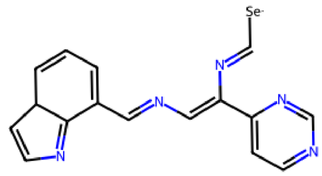

(d) Fitness: 8.91
$SA_{score}$: 5.09
Tanimoto: 0.11

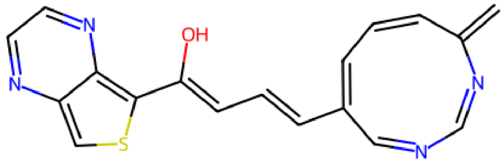

(e) Fitness: 9.14
$SA_{score}$: 4.68
Tanimoto: 0.14

Figure S3: **Generated molecules for the $PCE_{PCBM} - SA_{score}$ task.** Molecules generated by our model for the 5 independent runs on the $PCE_{PCBM} - SA_{score}$ maximization task.

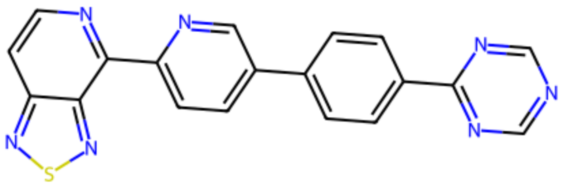

Figure S4: **Best dataset molecule for the $PCE_{PCBM} - SA_{score}$ task.** Molecule in the dataset with the highest fitness score for the $PCE_{PCBM} - SA_{score}$ maximization task, with fitness score 7.57 and $SA_{score}$ 4.59

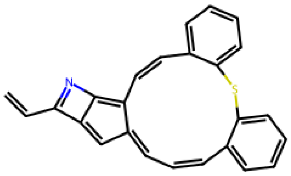

(a) Fitness: 28.29
$SA_{score}$: 2.90
Tanimoto: 0.18

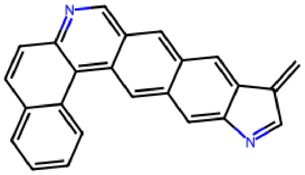

(b) Fitness: 26.95
$SA_{score}$: 2.90
Tanimoto: 0.53

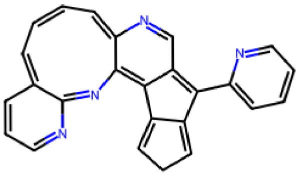

(c) Fitness: 28.65
$SA_{score}$: 4.35
Tanimoto: 0.17

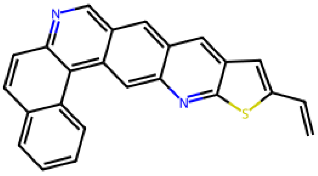

(d) Fitness: 27.09
$SA_{score}$: 2.85
Tanimoto: 0.55

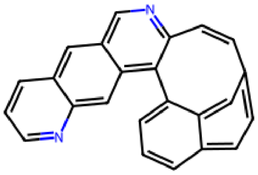

(e) Fitness: 29.22
$SA_{score}$: 3.52
Tanimoto: 0.47

Figure S5: **Generated molecules for the $PCE_{PCDTBT} - SA_{score}$ task.** Molecules generated by our model for the 5 independent runs on the $PCE_{PCDTBT} - SA_{score}$ maximization task.

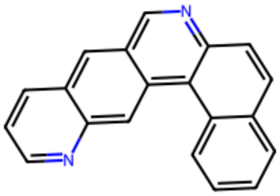

Figure S6: **Best dataset molecules for the $PCE_{PCDTBT} - SA_{score}$ task.** Molecule in the dataset with the highest fitness score for the $PCE_{PCDTBT} - SA_{score}$ maximization task, with fitness score 31.71 and $SA_{score}$ 2.20

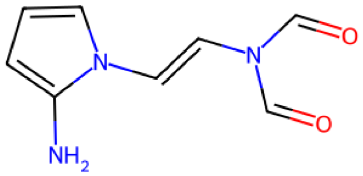

(a) Fitness: 0.0045
$SA_{score}$: 4.18
Tanimoto: 0.27

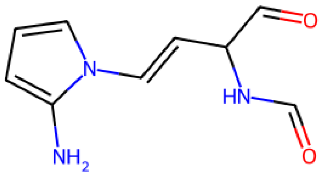

(b) Fitness: 0.0034
$SA_{score}$: 4.50
Tanimoto: 0.25

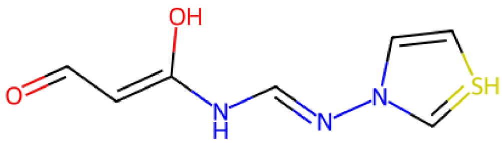

(c) Fitness: $7 \times 10^{-15}$
$SA_{score}$: 5.62
Tanimoto: 0.09

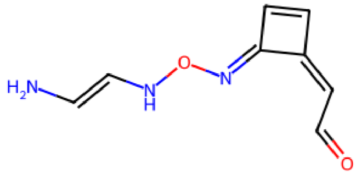

(d) Fitness: 0.0015
$SA_{score}$: 4.78
Tanimoto: 0.09

Figure S7: **Generated molecules for the singlet-triplet task.** Molecules generated by our model for the 5 independent runs on the singlet-triplet minimization task. Molecule **??** was proposed twice.

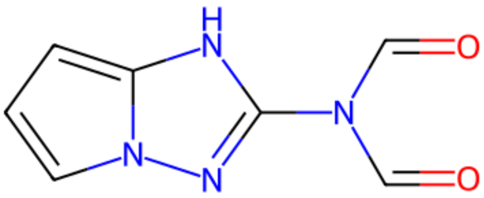

Figure S8: **Best dataset molecules for the singlet-triplet task.** Molecule in the dataset with the highest fitness score for the singlet-triplet task, with fitness score 0.020 and $SA_{score}$ 4.03

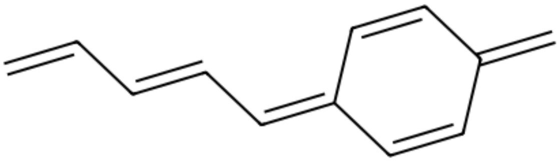

Figure S9: **Best dataset molecules for the oscillator strength task.** Molecule in the dataset with the highest fitness score for the oscillator strength maximization task, with fitness score 2.97 and $SA_{score}$ 3.79

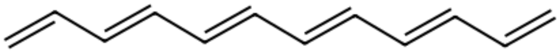

Figure S10: **Generated molecules for the oscillator strength task.** Molecule generated by our model in 3 out of 5 runs of the oscillator strength maximization task, with fitness 3.68, $SA_{score}$ 3.41 and a Tanimoto similarity of 0.33 with the initial molecule

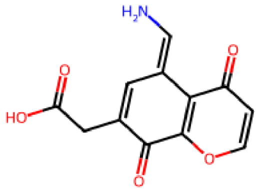


(a) Fitness: $-10.7$
$SA_{score}$: 3.45
Tanimoto: 0.26

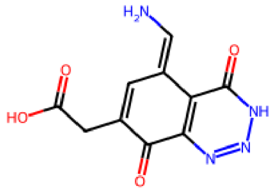


(b) Fitness: $-10.7$
$SA_{score}$: 3.75
Tanimoto: 0.26

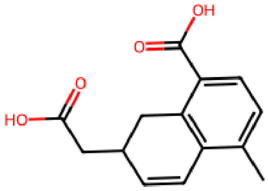


(c) Fitness: $-10.7$
$SA_{score}$: 3.33
Tanimoto: 0.26

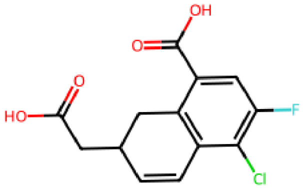


(d) Fitness: $-10.7$
$SA_{score}$: 3.54
Tanimoto: 0.23

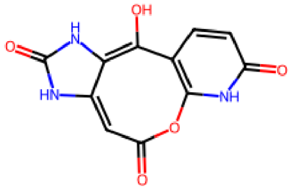


(e) Fitness: $-11.2$
$SA_{score}$: 3.78
Tanimoto: 0.25

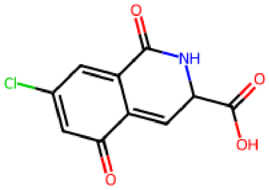


(f) Fitness: $-11.1$
$SA_{score}$: 1.90
Tanimoto: 0.18

Figure S11: **Generated molecules for the 1SYH task.** Molecules generated by our model in the 1SYH docking task

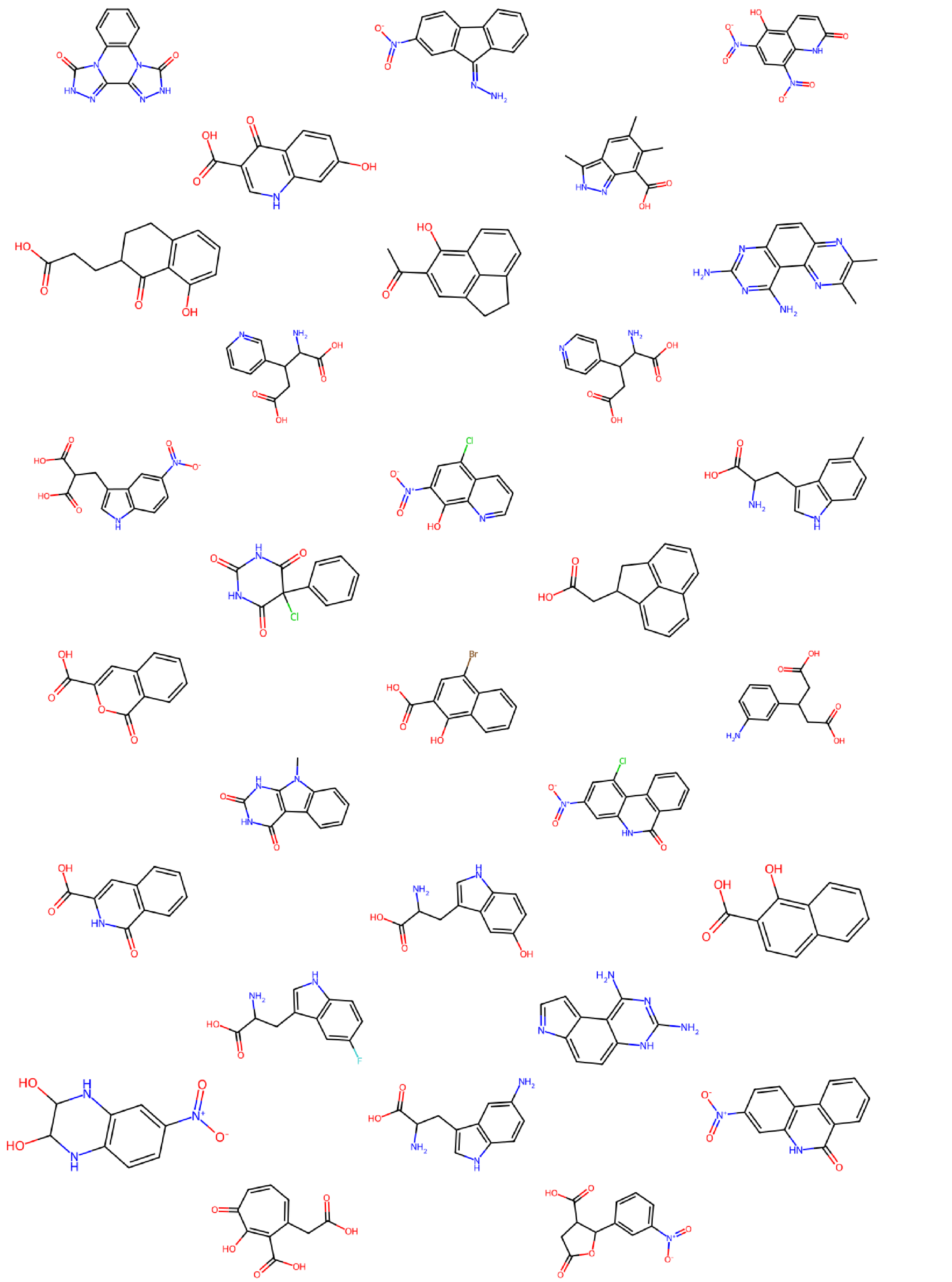

Figure S12: **Best dataset molecules for the 1SYH task.** All molecules with the best affinity score with 1SYH ($-9.9$) in the initial dataset

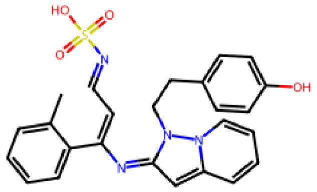

(a) Fitness: $-7.5$
$SA_{score}$: 3.57
Tanimoto: 0.14

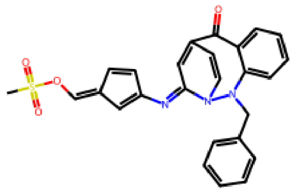

(b) Fitness: $-7.5$
$SA_{score}$: 4.29
Tanimoto: 0.21

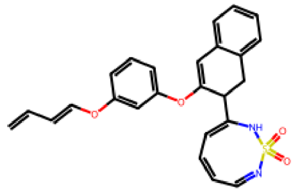

(c) Fitness: $-7.8$
$SA_{score}$: 4.28
Tanimoto: 0.09

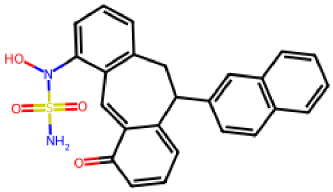

(d) Fitness: $-8.6$
$SA_{score}$: 3.68
Tanimoto: 0.12

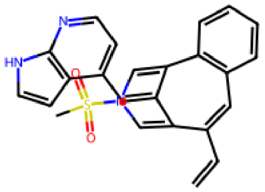

(e) Fitness: $-7.7$
$SA_{score}$: 2.30
Tanimoto: 0.12

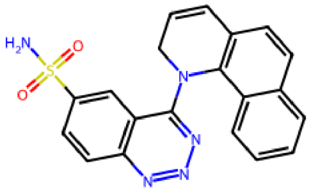

(f) Fitness: $-8.3$
$SA_{score}$: 2.89
Tanimoto: 0.27

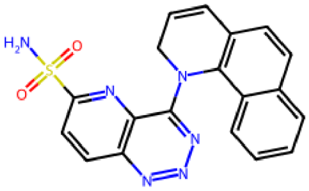

(g) Fitness: $-8.3$
$SA_{score}$: 2.30
Tanimoto: 0.20

Figure S13: **Generated molecules for the 6Y2F task**. Molecules generated by our model in the 6Y2F docking task

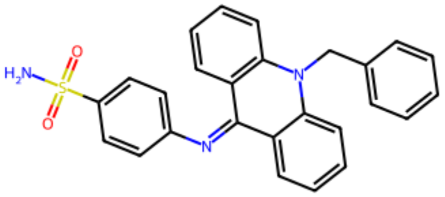

Figure S14: **Best molecule in the dataset for the 6Y2F task.** Molecule in the dataset with the best affinity with 6Y2F (Fitness: −8.2, $SA_{score}$: 2.30)

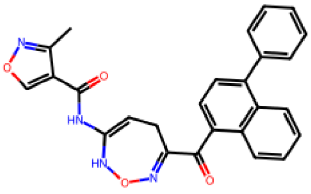

(a) Fitness: −13.6
$SA_{score}$: 3.16
Tanimoto: 0.29

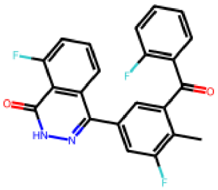

(b) Fitness: −14.6
$SA_{score}$: 2.53
Tanimoto: 0.34

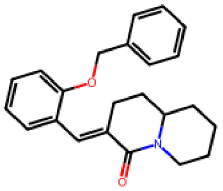

(c) Fitness: −12.3
$SA_{score}$: 2.59
Tanimoto: 0.36

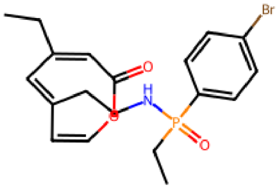

(d) Fitness: −11.9
$SA_{score}$: 4.30
Tanimoto: 0.20

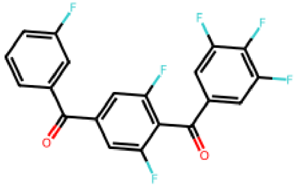

(e) Fitness: −13.3
$SA_{score}$: 2.35
Tanimoto: 0.37

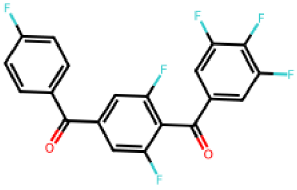

(f) Fitness: −13.3
$SA_{score}$: 2.27
Tanimoto: 0.39

Figure S15: **Generated molecules for the 4LDE task.** Molecules generated by our model in the 4LDE docking task
